# Cryo-EM structure of Slo1 with the auxiliary γ1 subunit suggests mechanism of depolarization-independent activation

**DOI:** 10.1101/2023.11.01.565197

**Authors:** Milena Redhardt, Stefan Raunser, Tobias Raisch

## Abstract

Mammalian Ca^2+^-dependent Slo K^+^ channels are expressed with β and γ auxiliary subunits that greatly influence voltage- and Ca^2+^-induced gating, thereby fundamentally altering the behavior of the channel. The four γ subunits reduce the need for voltage-dependent activation, allowing Slo to open in the absence of an action potential. The mechanism of this activation has, however, remained elusive. Here, we present the cryo-EM structure of Slo1 in complex with γ1/LRRC26, revealing how the transmembrane helix of γ1 binds and presumably stabilizes the active conformation of the voltage-sensor domain. This effect is further enhanced by a polybasic stretch on the intracellular side of the membrane which locally changes the charge gradient across the membrane. Sequence differences explain why the four γ subunits possess different activation efficiencies. Simultaneous binding of γ and the unrelated β subunits is structurally possible, as both binding sites do not overlap and the γ1 LRR domains are partially flexible. Thus, our data provide a possible explanation for Slo1 regulation by γ subunits, and furthermore suggest a novel mechanism of activation of voltage-gated ion channels by auxiliary subunits and add to the growing knowledge of their complex regulation.

## Introduction

Slo channels are Ca^2+^-gated K^+^ channels that are important for neuronal function, skeletal muscle contraction and maintaining smooth muscle tension in metazoa ^1^. Their characteristic feature is an extraordinarily high single-channel conductance that is, in excitable cells, initiated through synergistic activation by transmembrane voltage potential depolarization and increasing intracellular Ca^2+^ levels ^2^. Slo channels, also called big potassium (BK) or maxiK channels, are ubiquitously expressed in many excitable cells, such as neurons and muscle cells, as well as non-excitable cells, such as epithelial cells ^3–6^. They are required for re-establishing the resting potential after an action potential, the release of neurotransmitters at synapses as well as preventing constant membrane depolarization on the postsynaptic side ^4^. Due to these important neuronal functions, Slo channels are the targets of many natural neurotoxins and candidate targets for novel insecticides ^7^.

Slo channels belong to the tetrameric voltage-gated ion channel superfamily in which each monomer provides minimally six transmembrane helices to the transmembrane domain: four helices S1-S4 comprise the voltage-sensor domain (VSD) and the two helices S5 and S6 stabilize the central selectivity filter hairpin. In vertebrate Slo1 and Slo3 as well as invertebrate Slo, S1-S6 are preceded by an additional N-terminal S0 helix running in a roughly antiparallel way along S3 that is absent in Slo2 and other voltage-gated ion channels ^8–13^. In helix S4 of the VSD, three arginines function as gating charges, i.e. they sense via their positively charged guanidinium groups the transmembrane voltage potential and upon depolarization move further towards the extracellular side, dragging S4 and the S4-S5 linker with them, resulting in a rearrangement of the entire VSD. This rearrangement acts in concert with Ca^2+^-induced opening of the cytosolic gate composed of pore domain helices S6. In all Slo channels, helix S6 is connected by an extended linker to a large, domain-swapped cytosolic gating ring containing a total of eight Ca^2+^-binding sites. Ca^2+^-binding induces a large outwards-movement of the most N-terminal domain of the gating ring, straightening the preceding linker. The resulting force on the S6 helix of the pore domain opens the cytosolic gate and allows K^+^ ions to reach the selectivity filter ^8–11^. The Ca^2+^-induced motion of the gating ring primes the VSDs by stabilizing a conformation where the gating-charge arginine residues in S4 have already partially moved to the activated positions, and vice versa, the active conformation of the VSD assists Ca^2+^-gating through contacts between the S0-S1 and S4-S5 linkers of the VSD and the gating ring in the Ca^2+^-bound conformation ^9^.

Many native ion channels, such as voltage-gated sodium and potassium channels, include auxiliary subunits which bind to the transmembrane domain and modulate its conformational space and energy landscape ^14–17^. As such auxiliary subunits often have very specific tissue expression patterns, they allow the differential tuning of the activity of a ubiquitously expressed channel ^18^. Two families of auxiliary subunits of Slo termed β and γ, each comprising four members, have been identified to date which are notably absent in invertebrates ^1,3,18,19^. They are especially important for the action of Slo1 in non-excitable cells since they change the Ca^2+^- and voltage-gating behavior of Slo1 and allow the activation of the channel under conditions where the Slo1 core tetramer alone is inactive ^18^.

In smooth muscle, Slo1 exists exclusively in a complex with β1 which is crucial to control muscle tone by an increased Ca^2+^ sensitivity ^20,21^. The other three β subunits are expressed in other tissues including chromaffin cells (β2) and in the brain (β4). β subunits increase the Ca^2+^ sensitivity of Slo1, decrease the activation and deactivation kinetics and also alter the sensitivity towards pore-blocking spider toxins ^3,22^. They bind into the wide groove between two neighboring voltage-sensor domains of Slo1 in a 1:1 stoichiometry via their two transmembrane helices ^10^. The small interspersed extracellular domains of all four β4 molecules associate with each other to form a basket. The function of these extracellular domains is still largely unknown.

The first evidence of a second type of auxiliary subunit was the observation that the leucine-rich repeat containing protein 26 (LRRC26) induces a large −140 mV negative shift of voltage dependance of Slo1 in non-excitable prostate cancer cells ^23^. Three more members of this family with a similar architecture of one transmembrane helix and an extracellular leucine-rich repeat domain were identified later ^3,18,19,24^. γ1-4 are expressed in a wide range of tissues including smooth muscle (LRRC26/γ1), testis (LRRC52/γ2), brain (LRRC55/γ3), the adrenal gland (LRRC38/γ4) and skeletal muscle (γ2 and γ4) ^24–26^. They have been termed γ subunits as they differ not only in their molecular architecture from β subunits, but also in the effect they have on Slo1 (and Slo3 in the case of γ2). They all induce negative shifts to different extents in voltage dependence and it was shown for γ1 that it strongly enhances allosteric coupling of voltage sensing and pore opening, thus explaining how Slo1 can be activated in non-excitable cells in the absence of an action potential or strongly changing intracellular Ca^2+^ levels ^19,23,25,27,28^. On the other hand, γ subunits do not affect the Ca^2+^ sensitivity. γ and β subunits can even be present in the same Slo1 channel, increasing the possibilities for very fine tuning of K^+^ translocation even further ^29^.

While the LRR domains do not contribute to the shift in voltage dependence induced by γ subunits, it was shown that both the transmembrane helix and a directly adjacent short polybasic stretch on the C-terminal tails are the critical determinants for this shift, indicating a direct structural effect on the transmembrane domain of Slo. Furthermore, experiments with chimeric constructs have demonstrated that the transmembrane helices of γ1 and γ2 are very effective and contribute to voltage shifting much more than the transmembrane helices of γ3 and γ4. Conversely, the polybasic stretches of γ1 and γ3 are more potent in inducing voltage dependency shifts compared to γ2 and γ4 ^30^. However, it remained unclear how γ subunits bind to the transmembrane domain of Slo1 and what the molecular basis for the observed negative shifts in voltage dependence and the differences between the four γ subunits is.

Here, we present a cryo-EM structure of γ1 bound to a Slo1 tetramer. We show that the transmembrane helix of γ1 binds to the distal face of the Slo1 VSD by perfect shape complementarity and that this arrangement is stabilized by a kink in the γ1 transmembrane helix and probably by a covalent cystine bridge between γ1 and the VSD. Furthermore, in our structure, a polybasic stretch extends the γ1 transmembrane helix into the cytosol and we propose that this polybasic stretch, by introducing immobile positive charges, helps the VSD to adopt its activated conformation even at resting potential. Sequence comparison of these structural elements between the γ subunits helps to understand their different effectiveness in inducing voltage dependence sights in Slo1. Finally, we show that simultaneous binding of β and γ subunits is structurally possible under the condition that the extracellular LRR domain of γ subunits, which form a flexible tetramer in the case of γ1 in our structure, would open up and rearrange.

## Results and discussion

In order to gain structural and mechanistic insight into the regulation of Slo1 by auxiliary subunits, we co-expressed eGFP-Strep-tagged rabbit Slo1 with either of the four mCherry-His_10_-tagged γ subunits (Figure 1A, Supplementary Figure 1). We then purified the complex of Slo1 and γ1/LRRC26 via a double affinity protocol (Supplementary Figure 2A). The complex eluted from the size exclusion chromatography column in a single symmetric peak and SDS-PAGE confirmed its purity (Supplementary Figure 2B,C). Using cryo-EM and single-particle analysis, we then determined a C4-symmetrized reconstruction of the purified Slo1-γ1 complex at a resolution of 2.4 Å (Figure 1C, Supplementary Figure 3, Table 1) which allowed us to build a molecular model of almost full-length Slo1 and the transmembrane region (residues R251-Q298) of γ1 (Figure 1C,D, Supplementary Figure 3E). Furthermore, we observed a very diffuse density centrally above the selectivity filter (Supplementary Figure 5A), which we attribute to the leucine-rich repeat domains of γ1.

**Figure 1:**
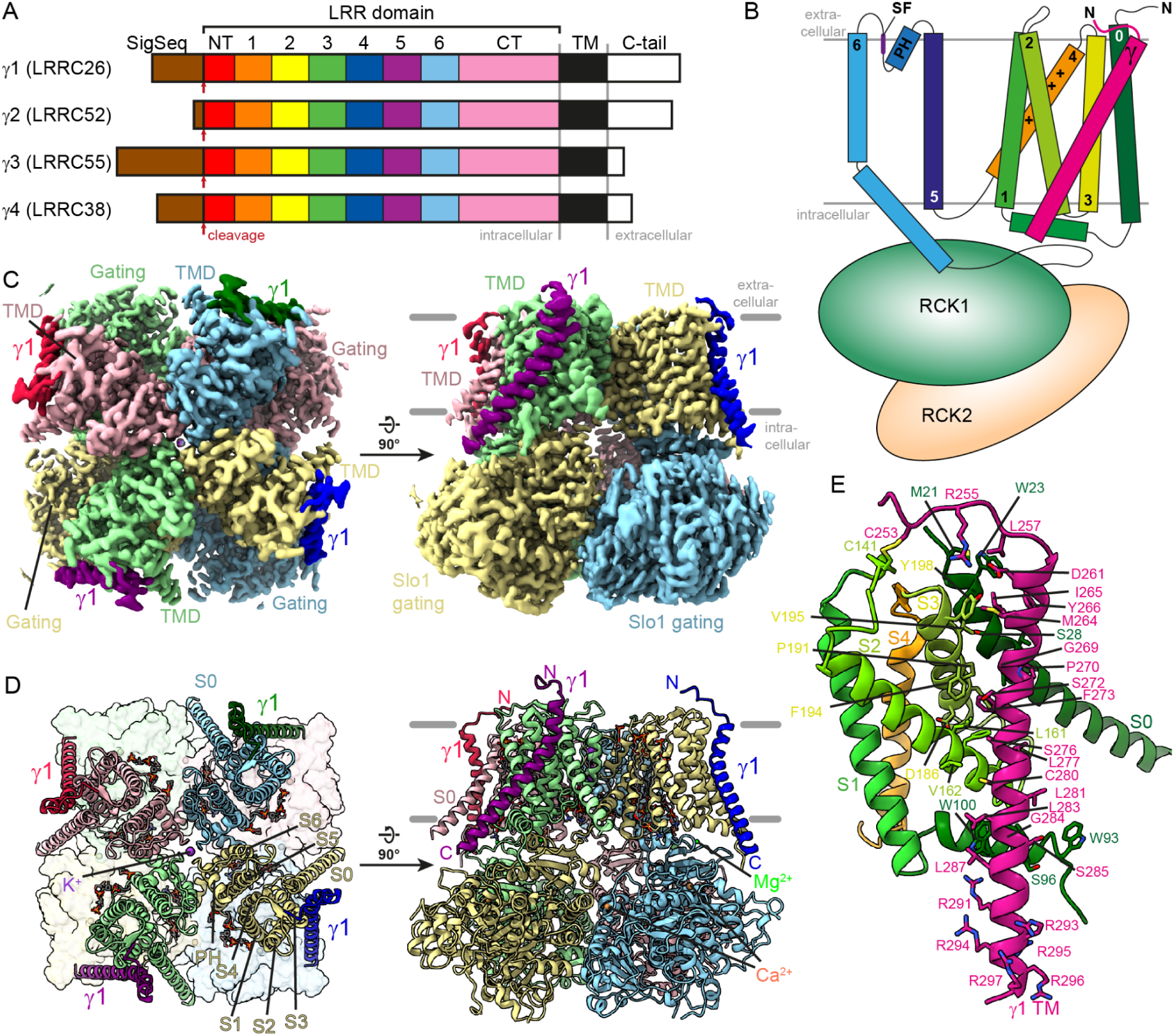
The transmembrane helix of γ1 binds to the voltage-sensor domain of Slo1. A) Schematic representation of γ subunits. γ1 - γ4 possess highly similar architectures comprising an N-terminal membrane-targeting signal sequence, an extracellular domain containing eight leucine-rich repeats, a transmembrane domain and a short C-terminal cytosolic tail. B) Schematic of the Slo1 transmembrane domain topology and the γ1 transmembrane helix which binds diagonally across helices S0, S3, S2 and the perpendicular S0-S1 connector helix, as observed in our Slo1-γ1 structure. C) Cryo-EM reconstruction of rabbit Slo1 in complex with γ1 in two orientations. Clear density corresponding to the γ1 transmembrane helix is visible at the four voltage-sensor domains. D) Model of the γ1-bound rabbit Slo1 in the same orientations as in panel C. The γ1 TM helices bind to the four corners of the transmembrane domain tetramer, contacting the voltage-sensor domains. E) The γ1 TM helix forms extensive hydrophobic contacts with helices S0, S0-S1, S2 and S3 of the Slo1 voltage-sensor domain and binds with perfect shape complementarity.

**Table 1:**
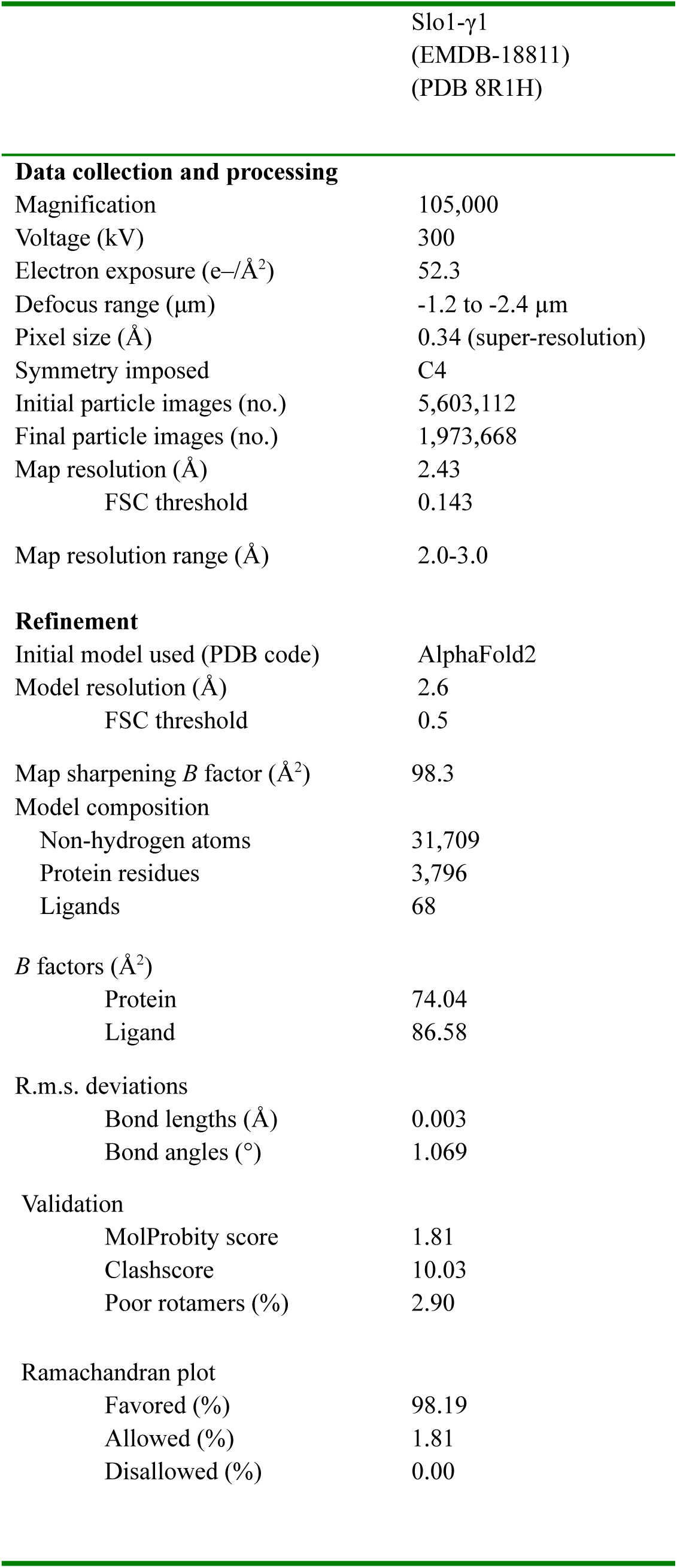
Cryo-EM data collection, refinement and validation statistics.

### Overall structure of the Slo1-γ1 complex

Rabbit Slo1 adopts a very similar overall structure and conformation to the previously published cryo-EM structures of Ca^2+^-bound human Slo1 (PDB 6V38)^10^, *Aplysia* Slo1 (PDB 5TJI)^9,11^ and *Drosophila* Slo (PDB 7PXE)^8^, with RMSDs of 0.75 Å, 3.07 Å and 1.79 Å, respectively (Supplementary Figure 4A-C). The transmembrane module of Slo1 harbors the central selectivity filter in which four K^+^ ions are stabilized by the conserved TVGYGD motif and the pore domain helices S5 and S6. This pore domain is surrounded by four voltage-sensor domains constructed of helices S0-S4 that extend to the four corners of a square when looking at the membrane from the extracellular side (Figure 1B,D). Helix S6 connects via an extended linker to the first of two ‘regulator of potassium conductance’ (RCK) domains. The eight RCK domains of the Slo tetramer associate into a large but compact gating ring that makes up the majority of the mass of the channel. In agreement with the presence of 10 mM Ca^2+^ during purification, all eight Ca^2+^-binding sites in the gating ring are occupied by ions and Slo1 adopts the active conformation characterized by an expanded gating ring, an extended S6 linker and an open intracellular gate ^8,10,11^. Similarly, the previously described Mg^2+^-binding sites between the gating ring and the VSDs are occupied^10,11^.

### γ1 binds to the distal face of the voltage-sensor domains of Slo1

The core of the interaction with Slo1 is mediated by the extended, kinked transmembrane helix of γ1 (residues P259-C290) that binds along the distal side of the VSD (Figure 1B-E). The interaction is predominantly hydrophobic and driven by perfect shape complementarity between γ1 and the Ca^2+^-activated conformation of the VSD. The most prominent feature of the γ1 helix is a kink at the outer leaflet of the lipid bilayer orchestrated by G269 which is invariable between γ subunits and the subsequent P270 that is conserved only among some γ1 homologs (Figure 1E and 2A-C, Supplementary Figure 1). The conformation of that kink running through a cleft along S0, S3 and S2 of the VSD is further stabilized by the flanking Y266 that caps the interaction towards the outside of S0 and F273 that inserts by hydrophobic and *π*-stacking interactions into a pocket spanned by helices S3 and S2 (Figure 1 E and 2C). On the intracellular edge of the membrane, the S0-S1 connector helix which runs roughly parallel to the membrane plane serves as a lock for the C-terminal part of the γ1 helix, effectively clamping it between its tryptophan residues W93 and W100, the latter touching the highly conserved G284 on γ1 (which is an alanine in γ4) (Figure 1E). The γ1 helix then continues at least three more turns into the cytosol until it almost touches the gating ring (Figure 1 C,D).

**Figure 2:**
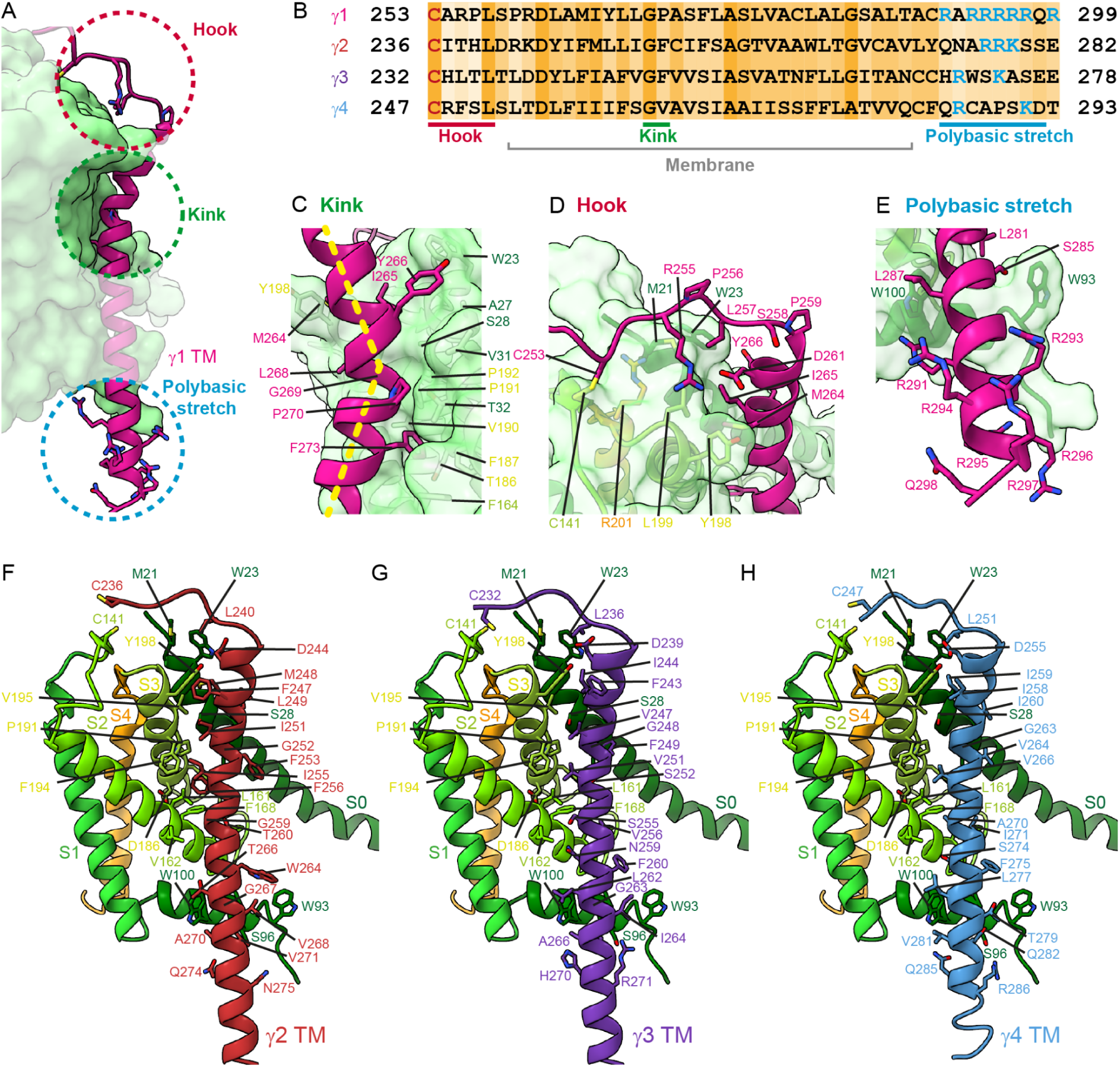
Three structural elements of γ subunits modulate the function of Slo1. A) Three structural elements in γ1 likely contribute to VSD activation: a kink in the central transmembrane helix, a hook N-terminal of it and a polybasic stretch on the intracellular side. B) Alignment of rabbit γ subunit transmembrane region containing the hook, the kink and the polybasic stretch. The cysteine in the hook which potentially forms a disulfide bond with Slo1 C141 is marked in red and the basic residues in the polybasic stretch in blue. C) Close-up of the kink with important residues on both sides shown in stick representation. The half-transparent surface of the VSD illustrates the shape-complementarity with γ1. D) Close-up of the hook region of γ1 N-terminal of the transmembrane helix. The hook contacts the VSD via R255, the invariant L257 and the proposed cystine bridge of C253. E) Close-up of the helical extension of the transmembrane helix that protrudes into the cytosol and harbors the polybasic stretch. In rabbit γ1, the polybasic stretch contains seven arginine residues and thereby produces a strongly positively charged microenvironment close to the voltage-sensor domain. F) Structure prediction of γ2 binding to the Slo1 VSD. The models in panels F)-H) were created by AlphaFold-Multimer, followed by careful manual adjustments. G) Structure prediction of γ3 binding to the Slo1 VSD. H) Structure prediction of γ4 binding to the Slo1 VSD.

It was previously reported based on conductance measurements using chimeric γ subunit constructs that the transmembrane helices and the C-terminal cytosolic tails are critical determinants of the observed differences in the ability of the γ subunits to negatively shift the Slo1 voltage dependence (γ1 > γ2 > γ3 > γ4). Furthermore, when comparing the transmembrane helices, γ1 and γ2 are much more effective compared to γ3 and γ4 ^30^. While sequence conservation of key residues and structure predictions using AlphaFold-Multimer ^31^ suggest that γ2, γ3 and γ4 bind Slo1 in a very similar manner (Figure 2B,F-H), sequence differences in several residues important for the interaction as well as slight differences in the AlphaFold-Multimer predictions (Figure 2B,F-H, Supplementary Figure 1) provide an explanation for the reported difference in effectiveness between the γ subunit transmembrane helices. Specifically, the phenylalanine at the very center of the interaction with the VSD (F273 in γ1 and F256 in γ2) is a serine in γ3 and γ4 (Figure 2B, Supplementary Figure 1), which weakens the overall interaction. Furthermore, P270 of γ1 which is involved in kink stabilization, is a phenylalanine in γ2 and γ3 and an aliphatic residue in γ4 (Figure 2B, Supplementary Figure 1), and those larger side chains possibly force the γ transmembrane helix in a different trajectory or lead to a locally different conformation of the VSD.

### γ1 presumably stabilizes the open conformation of Slo1

It has been observed that the −140 mV negative shift induced by γ1 does not change the Ca^2+^ sensitivity of Slo1 ^23^, and is described in the Horrigan-Aldrich model of Slo1 activation by 20-fold increase in the allosteric coupling of VSD activation and pore domain opening ^2,23^. On the structural level, depolarization of the membrane potential, i.e. a reversal of the local charge gradient across the membrane, leads to a dragging force on the positive arginine gating charges in the S4 helices of ion channels towards the extracellular side ^18,32^. This repositioning of the entire S4 is compensated by a structural change of the whole VSD module and the S4-S5 linker, which in turn primes Slo1 activation by allowing the S6 helix in the cytosolic gate to adopt the open conformation. Hence, a possible activation mechanism of γ1 might be to change the energy landscape for gating charge movement towards the extracellular side, which would make a VSD rearrangement and gate opening easier and more likely.

We have identified three structural elements which might act together to cause such a shift. The first is the above-mentioned shape complementarity of the γ1 transmembrane helix with respect to the surface of the partially activated state of the VSD (Figure 1E and 2A). In particular, the kink in the helix seems important as the helix must change direction to be accommodated at the concave surface of the VSD (Figure 2C). The perfect fit of γ1 to the partially activated conformation of the VSD might indicate a preference for this state. Conversely, preferential binding of γ subunits to this partially activated over the resting state might lower the energy cost of the voltage-driven rearrangement and thereby stabilize the active conformation. The kink is likely most pronounced in γ1 due to the G269-P270 amino acid tandem, and the absence of a proline in the equivalent position, together with the substitution of the central phenylalanine by serine, could be responsible for the lower effectiveness of activation by γ3 and γ4 transmembrane helices ^30^.

The second element is a hook comprising residues R251-P259 directly N-terminal of the transmembrane helix (Figure 2A,D). This hook binds along the extracellular side of the VSD, thereby stabilizing the interaction of the transmembrane helix. The direction of the hook is defined by P259 and the invariable L257 might play an important role as well (Figure 2B,D). However, the most interesting feature is C253 which is conserved in all γ subunits. The side chain of C253 is positioned very close to C141 of the VSD and we thus propose that Slo1 and γ1 might be connected via a cystine bond (Figure 2D), even though the limited local resolution does not allow a certain conclusion. Such covalent anchoring would directly strengthen the interaction of the kinked transmembrane domain and thereby help to stabilize the active conformation of the VSD. Cystine anchoring is known from the β2 and β4 subunits of mammalian voltage-gated sodium channels which also bind to the distal side of the VSD with the help of a single transmembrane helix and a disulfide bond and which also induce a (small) negative shift in voltage dependence ^33,34^.

Lastly, the γ1 helix extends several turns beyond the membrane on the intracellular side. In rabbit γ1, this R291-Q298 extension contains an impressive number of seven arginine residues (Figure 3A,B,E). It was shown before that this stretch is crucial for the voltage-dependence shift ^30^ and the positioning of this stretch directly besides the VSD suggests a compelling explanation for this observation. During resting potential, the intracellular side of the plasma membrane is negatively charged, while the extracellular side is positively charged. During depolarization, this is reversed, causing the gating charges in S4 to follow the voltage change and move towards the extracellular side ^32^. The presence of several immobile positive charges on the intracellular side in close proximity to the VSD as in the case of the Slo1-γ1 complex is likely to locally lower the resting state potential and repulse the gating charges, thereby reducing the energy to overcome for the VSD to transition to the active conformation.

**Figure 3:**
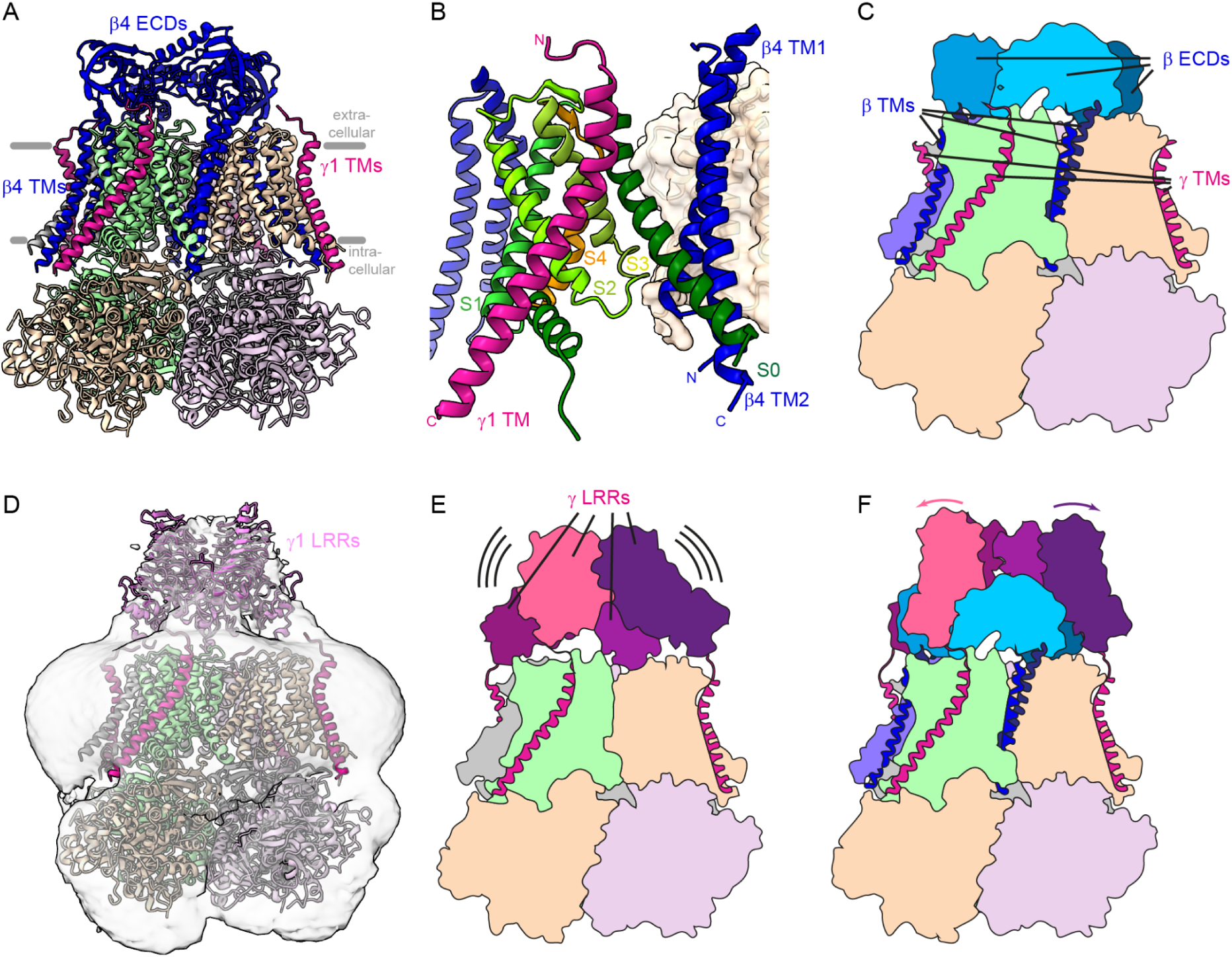
Simultaneous binding of β and γ subunits to Slo1 is structurally possible, but would require rearrangements in their extracellular modules. A) Superposition of the human Slo1-β4 structure (PDB 6V22) onto our rabbit Slo1-γ4 structure; the human Slo1 was omitted for clarity. The extracellular domains (ECDs) of the β subunits form a cage above the Slo1 transmembrane domain. B) Close-up view on one of the VSDs of Slo1 with bound β and γ subunit transmembrane helices as shown in panel A). While the γ TM helix binds diagonally at the most distal side of the VSD, one β transmembrane helix pair contacts the intracellular part of helix S0, while another β subunit contacts the VSD from the other side mostly via the extracellular S1-S2 loop. C) Cartoon model of Slo1 with bound β subunits and γ subunit transmembrane helices. D) At low threshold, weak density is visible centrally above the Slo1 transmembrane domains in a lower-resolution reconstruction (see also Supplementary Figure 5A). An AlphaFold-multimer model of a γ1 LRR tetramer (arrangement B in Supplementary Figure 5B) fits well into the density; distance between the C-terminus of the LRR and the N-terminus of the TM helix allows this configuration. E) Cartoon model of Slo1 with full-length γ subunits. The LRRs display a high degree of flexibility. F) Cartoon of Slo1 with bound β subunits and full-length γ subunits. A rearrangement (outwards movement) of γ LRRs would be necessary to prevent clashes with β ECDs. This would be the case for all arrangements shown in Supplementary Fig. 5B.

Interestingly, the number of positively charged residues in the polybasic stretch differs between different γ subunits proportional to their capability to induce negative shifts of the voltage dependence. It is highest in mammalian γ1 with six or seven, while γ2 contains three and γ3 and γ4 only two arginines or lysines (Figure 2B, Supplementary Figure 1). Accordingly, the C-terminal tails of γ1 and γ3 are much more effective in shifting the voltage dependence of Slo1 compared to γ2 and γ4 and this depends for all but γ2 entirely on the presence of the polybasic stretch ^30^.

Taken together, we propose that the combination of three different principles, namely shape complementarity, covalent anchoring and lowering the resting state potential by a positively charged intracellular stretch, act in concert to stabilize an active VSD conformation in the Slo1-γ1 complex. This mechanism would further comprehensively explain the different strengths of induced shifts in voltage dependence by the different γ subunits.

### Simultaneous binding of β and γ subunits

Earlier observations in electrophysiology experiments suggested that β and γ subunits can exist in the same Slo1 channels. More specifically, overexpression of Slo1 with β2 and γ1 resulted in a −120 mV shift in absence of Ca^2+^ with respect to Slo1 apo, which is a characteristic of γ1 regulation, while at the same time, a β2-specific complete inactivation occurs after activation ^29^. This observation is very much in line with the simultaneous presence of β2 and γ1 and prompted us to ask whether simultaneous binding is also structurally possible and whether rearrangements would be necessary.

We generated a composite model by superposing the β4-bound human Slo1 (PDB 6V22)^10^ onto our structure of γ1-bound rabbit Slo1. Indeed, both types of regulatory subunits bind the transmembrane domain of Slo1 using completely different and non-overlapping binding sites (Figure 3A-C). While γ1 binds to the distal side of the VSD with contacts to S0, S0-S1, S2 and S3 (Figure 1E and 3B), the two transmembrane domains of β4 fit in the cleft between two neighboring VSDs (Figure 3A), and contacts include the cytosolic part of S0 of one and the extracellular S1-S2 loop of the other VSD (Figure 3B); also lipid molecules mediate part of the interaction ^10^. The four ECDs of β4 are positioned at a short distance above the transmembrane helices in a cage-like tetrameric arrangement shielding the extracellular side of the Slo1 selectivity filter (Figure 3A,C). No clashes between the TM helices of β4 and γ1 are visible and the Slo1 VSD conformation is, apart from a minor shift of the anyway slightly mobile cytosolic part of S0, virtually identical between both structures (RMSD 0.825 Å over 257 Cα atoms of the transmembrane domain of a single chain). Thus, simultaneous binding of both types of regulatory subunits is possible at least in the transmembrane portions when not taking into account the extracellular domains.

### The leucine-rich repeat domains of γ1 constitute a flexible extracellular module

The role of the LRR domains of γ subunits has so far remained elusive. As shown with the help of chimeric γ constructs, only the transmembrane part and a short patch on the intracellular side determined the extent of the negative shift in voltage-dependence of Slo1 ^30^. On the other hand, the LRRs are the most conserved part of γ1-4 (Supplementary Figure 1), suggesting an important role for the function of the proteins. As the extracellular domains of β4 form a stable, albeit slightly mobile, cage that shields the extracellular side of the pore to some extent (Figure 3A,C)^10^, one aim of our study was to determine whether the γ1 LRRs might potentially form a similar stable, tetrameric arrangement.

In our 2D classes and the reconstructed EM volumes, we observe a very weak and diffuse density centrally above the transmembrane domains of Slo1 (Figure 3D and Supplementary Figure 3A and 5A). This density is absent in all of the Slo reconstructions in absence of an auxiliary subunit ^8–11^. We therefore attribute it to the four LRRs. The position of the LRRs is similar to the place where the β4 cage is localized in the structure of β4-bound human Slo1 (Figure 3D,E) ^10^. However, judging from the weakness of the density, the LRRs of γ1 seem to be much more flexible than the equivalent domains of the β4 cage.

AlphaFold-Multimer ^31^ reproducibly predicts the LRR domains of all four γ subunits as tetramers, but the arrangements differ from each other (Supplementary Figure 5B). Several such arrangements are sterically possible in the context of a Slo1-γ complex because all four transmembrane helices point in the same direction and the spacing of the C-terminal ends of the LRRs is compatible with the position of the transmembrane helices at the four corners of the Slo1 transmembrane domain square (Figure 3D,E). When positioned on the N-terminal ends of the transmembrane helices in our cryo-EM structure, the AlphaFold model of the LRRs of γ1 would indeed be cage-like and fit in the density above the center of the Slo1 transmembrane domain (Figure 3D,E). Such a cage-like structure would also explain the observed slight decrease in efficiency of Slo1 inhibition by pore-blocking toxins in presence of γ1 ^28^, as it would pose a kinetic barrier for toxin binding. Thus, with regard to our data and in context of the literature, we propose that the LRR domains of γ subunits might assemble in a cage-like, mobile arrangement. However, the main physiological role of the LRRs still needs to be determined and might include binding to extracellular matrix proteins as has been suggested previously for the extracellular domains of β subunits ^10^. A further question which arises from our structure is how the ECDs of β subunits and the LRRs of γ subunits would arrange in case they are present on the same Slo1 tetramer, as the arrangement of LRRs we observe for γ1 would clash with the cage formed by a β subunit tetramer. Since the γ1 LRR domains are relatively flexible, we propose that the LRR tetramer might open up in this case to prevent clashes with the β ECDs (Figure 3F).

## Conclusion

In this study, we report the cryo-EM structure of a Slo1-γ1 complex and propose a mechanism how γ1 might shift the voltage dependence of the channel to values that allow activation in non-excitable cells under physiological conditions, i.e. in the absence of an action potential-associated membrane depolarization (Figure 4). The γ1 transmembrane helix itself contains a prominent kink which allows it to perfectly fit along the concave surface of the VSD in the active conformation. This association is stabilized by a hook on the extracellular side that might involve a cystine bridge. And finally, a polybasic stretch introduces immobile positive charges on the intracellular side of the membrane, which might locally decrease the resting potential and prime the VSD for activation (Figure 4). Differences in the sequences of the transmembrane helices and the polybasic stretches could explain the different potencies of the four γ subunits in VSD activation. This proposed model is in line with previous functional work that has identified the polybasic stretch as a critical determinant for γ subunit activity ^30^. Our structural work thereby provides a probable explanation for Slo1 activation in non-excitable cells by γ subunits.

**Figure 4.**
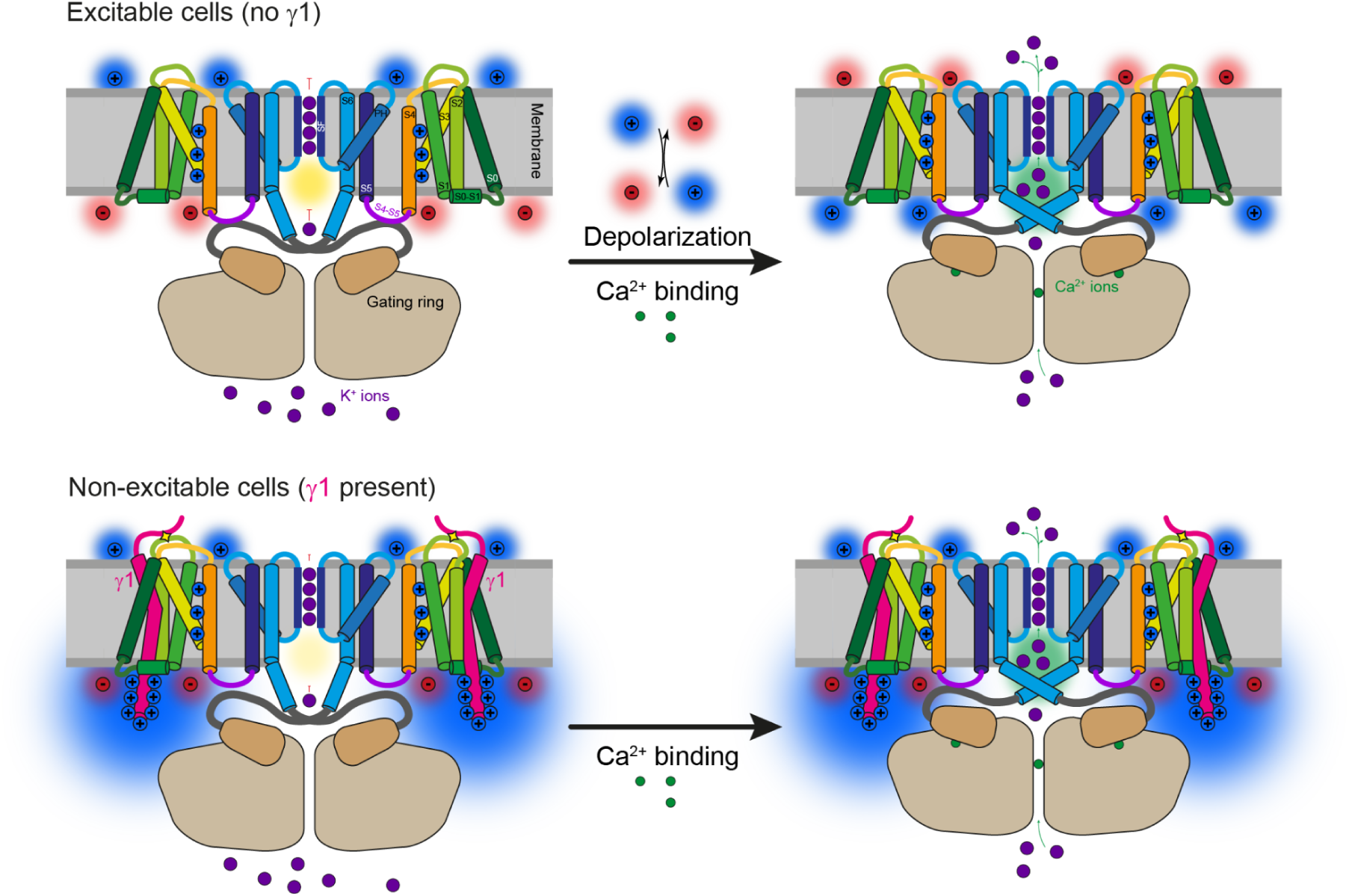
Proposed model for Slo1 activation by γ1. In the absence of γ subunits, Slo1 is activated in excitable cells synergistically by membrane potential depolarization and intracellular calcium binding. In the resting state, a charge gradient over the membrane exists with negative net charges on the intracellular side and positive charges on the extracellular side. This distribution favors the resting state conformation of the voltage-sensor domains of Slo1 in which the three (positively charged) arginine gating charges reside close to the intracellular side of the membrane. Depolarization, i.e. charge gradient reversal, pulls the gating charges towards the extracellular side, leading to a rearrangement throughout the entire VSD. This is synergistically enhanced by binding of calcium ions to the gating ring, which induces an outwards movement of the S6 helix and opening of the intracellular gate. In non-excitable cells, where the resting state transmembrane potential is constant, we propose that the positive charges in the C-terminal region of the γ subunits locally change the membrane potential and induce a similar movement of the Slo1 gating charges towards the extracellular side as observed during action potential. Thus, the need for the voltage depolarization signal is decreased or completely absent and only binding of calcium is necessary to open Slo1.

Beyond this, it will be interesting to experimentally test our hypothesis about simultaneous binding of β and γ subunits, especially regarding the necessary rearrangements within the extracellular domains (Figure 3F). Also, future studies should investigate in more detail the molecular roles of the β and γ extracellular domains which, due to their high conservation, must certainly be important for Slo1 function.

Besides Slo, also many other ion channels including voltage-gated sodium, potassium and calcium channels as well as TRP channels contain auxiliary subunits which regulate their activation ^14,16,17^. While cystine bond anchoring has been shown at least for Na_V_ β2 and β4 ^33,34^, other auxiliary subunits which induce local changes in transmembrane voltage potential by placing immobile charges close to the VSD have not been reported to the best of our knowledge. Thus, the activation mode by a polybasic stretch as we propose for Slo γ subunits is a novel mechanism that expands the repertoire of auxiliary subunits, and it will be very interesting to see whether similar modes of action are also applied by other proteins.

## Material and Methods

### Cloning and expression

*cDNA* encoding rabbit Slo1 (*KCNMA1*; Uniprot Q9BG98, residues 1-1118), was kindly provided by Mark Hollywood (Dundalk Institute of Technology) ^35^ and was subcloned into an altered pEG BacMam vector ^36^. The fusion construct carries a C-terminal, HRV 3C-cleavable eGFP-2xStrep tag.

cDNAs encoding the four rabbit γ subunits were ordered as synthetic, codon-optimized DNA fragments (Thermo Fisher Scientific) and also cloned into pEG BacMam vectors. The resulting fusion proteins carried HRV 3C-cleavable mCherry-His_10_ tags.

Heterologous expression of Slo1 and γ subunits in HEK293S GnTI-cells was performed using the BacMam method ^36^. Bacmids containing either Slo1 or γ1-4 cDNA were produced in *E. coli EMBacY* cells by transforming the respective pEG BacMam constructs.

*Spodoptera frugiperda* Sf9 cells were then transfected with these bacmids using FuGene Transfection Reactant (Promega). Baculoviruses were amplified in Sf9 cells at 28 °C.

Slo1 and γ subunits were co-expressed in HEK293S GnTI-cells. Cells were cultured at 37 °C to a density of ∼ 3×10^6^ cells/ml and infected with 7% (v:v) Slo1 baculovirus and 11% (v:v) γ1 baculovirus. After 8 h, 10 mM sodium butyrate was added, and the temperature was reduced to 30°C. ∼ 44 h post-induction, cells were collected by centrifugation and stored at −80°C until further use.

### Purification of the Slo1-γ1 complex

Purification of the Slo1-γ1 complex was performed according to ^8,11^.

All purification steps were performed either on ice or at 4 °C. Cells were lysed by homogenization in hypotonic buffer containing 10 mM Tris-HCl pH 8.0 and 2 mM EDTA, supplemented with protease inhibitors (cOmplete ™ Mini EDTA-free Protease Inhibitor Cocktail; Roche). The cell lysate was centrifuged for 15 min at 38,000 g and the membrane fraction was resuspended in hypotonic buffer. The lysate was centrifuged again for 15 min at 38,000 g and the membrane fraction was then resuspended in a basic buffer containing 20 mM Tris-HCl pH 7.6, 320 mM KCl, 10 mM CaCl_2_ and 10 mM MgCl_2_, supplemented with protease inhibitors. Membranes were solubilized for 3 h using 1% (w/v) lauryl maltose neopentyl glycol (LMNG) (Neo Biotech) and 0.1% (w/v) cholesteryl hemisuccinate (CHS) (Sigma Aldrich). Subsequently, the sample was centrifuged at 38,000 g for 30 min.

The solubilized fraction was incubated with Strep-Tactin beads (IBA Lifesciences) for 1 h. Then, the supernatant was removed and the beads washed with basic buffer supplemented with 0.05 % LMNG and 0.005 % CHS. Then, the Slo1-γ1 complex was eluted using basic buffer supplemented with 0.05 % LMNG, 0.005 % CHS, 20 mM imidazole and 2.5 mM d-desthiobiotin.

The eluate of the Strep-Tactin beads was applied to Ni-NTA beads (QIAGEN). After 1 h, the supernatant was removed and beads washed with basic buffer supplemented with 20 mM imidazole, 0.05 % LMNG and 0.005 % CHS. The Slo1-γ1 complex was eluted by a step gradient to basic buffer supplemented with 300 mM imidazole, 0.05 % LMNG and 0.005 % CHS. Finally, the Ni-NTA eluate was applied to size exclusion chromatography on a Superose 6 Increase 5/150 column (GE Healthcare) in basic buffer supplemented with 0.003 % LMNG and 0.0003 % CHS.

The peak fractions corresponding to the Slo1-γ1 complex were concentrated to 4.3 mg/ml and used for cryo-EM grid preparation.

### Cryo-EM sample preparation and data acquisition

Grids were prepared using a Vitrobot Mark IV (Thermo Fisher Scientific) at 13°C and 100% humidity. 4 μl of 4.3 mg/ml Slo1-γ1 complex were applied to glow-discharged UltrAuFoil R2/2 200 grid (Quantifoil). Any excess liquid was removed by blotting for 3.5 s at a blot force of −3, before the grids were vitrified by plunging them into liquid ethane. cryo-EM data were acquired on a Cs-corrected Titan Krios G3 electron microscope (Thermo Fisher Scientific) equipped with a field emission gun. With a K3 camera (Gatan), a total of 16,802 movies were recorded in super-resolution mode at a nominal magnification of 105,000 x. This resulted in a super-resolution pixel size of 0.34 Å. Zero-loss filtration was performed using a Bioquantum post-column energy filter (Gatan), with a slit width of 15 eV. Distributed over 60 frames, the total electron exposure was 52.3 e^-^/Å^2^.

The dataset was collected using the automated data acquisition software EPU (Thermo Fisher Scientific). Four acquisitions were acquired per hole using a defocus range of −1.2 to −2.4 µm. Details and statistics of data acquisition can be found in Table 1.

### Data processing and model building

Data preprocessing in tranSPHIRE ^37^ included motion correction by MotionCor2 ^38^ and CTF estimation in CTFFIND4 ^39^. 5,603,112 particles were picked using SPHIRE-crYOLO ^40^ and extracted using a box size of 256×256 pixels after 2-fold binning. 2D classification was performed within cryoSPARC ^41^. 3,284,471 particles that were assigned to well resolved classes were used to create three *ab initio* models that were then used as references in a heterogeneous refinement; this and all further reconstruction steps were performed while applying C4 symmetry. A subsequent homogeneous refinement using the 2,196,730 particles assigned to the best class resulted in an initial 2.9 Å reconstruction. These particles were then converted and further cleaned by 3D classification in RELION 3.1 ^42^. 1,973,668 particles were subjected to particle polishing in RELION followed by non-uniform refinement including CTF refinement in cryoSPARC, yielding a resolution of 2.43 Å. The final reconstruction was post-processed in PHENIX ^43^ by applying a sharpening B-factor of 98.3 Å^2^. The density map shown in Figure 1C has been post-processed using DeepEMhancer ^44^.

An initial molecular model of a Slo1 tetramer with bound γ1 transmembrane and cytosolic region (residues 250-329) was created using AlphaFold-Multimer ^31^, manually truncated and adjusted in COOT ^45^ and optimized using real space refinement against the PHENIX-sharpened real space map in PHENIX ^43^. Refinement restraints for the LMNG molecule were created using PHENIX.Elbow. All structural figures were created using UCSF ChimeraX ^46^.

Details and statistics of data processing and model building can be found in Supplementary Figure 3 and Table 1.

## Acknowledgements

We thank Nathalie Bleimling and Marion Hülseweh for technical support in the wet lab, and Oliver Hofnagel and Daniel Prumbaum for assistance with recording cryo-EM data. This work was supported by the Max Planck Society (to S.R.) and a grant of the Deutsche Forschungsgemeinschaft (German Research Foundation) TRR 152 (project-ID 239283807 to S.R.).

## Competing interests

The authors declare no competing interests.

## Data availability

The cryo-EM data have been deposited at the EMDB under the accession number EMD-18811 and the molecular model at the PDB under the accession number 8R1H.

## Author contributions

T.R. conceived the study. S.R. acquired funding. M.R. cloned, expressed and purified the Slo1-γ1 complex and prepared cryo-EM samples. M.R. and T.R. acquired cryo-EM data. T.R. processed the data and built the molecular model. All authors analyzed data. T.R. wrote the manuscript with input from the other authors.

---

**Supplementary Figure 1:**
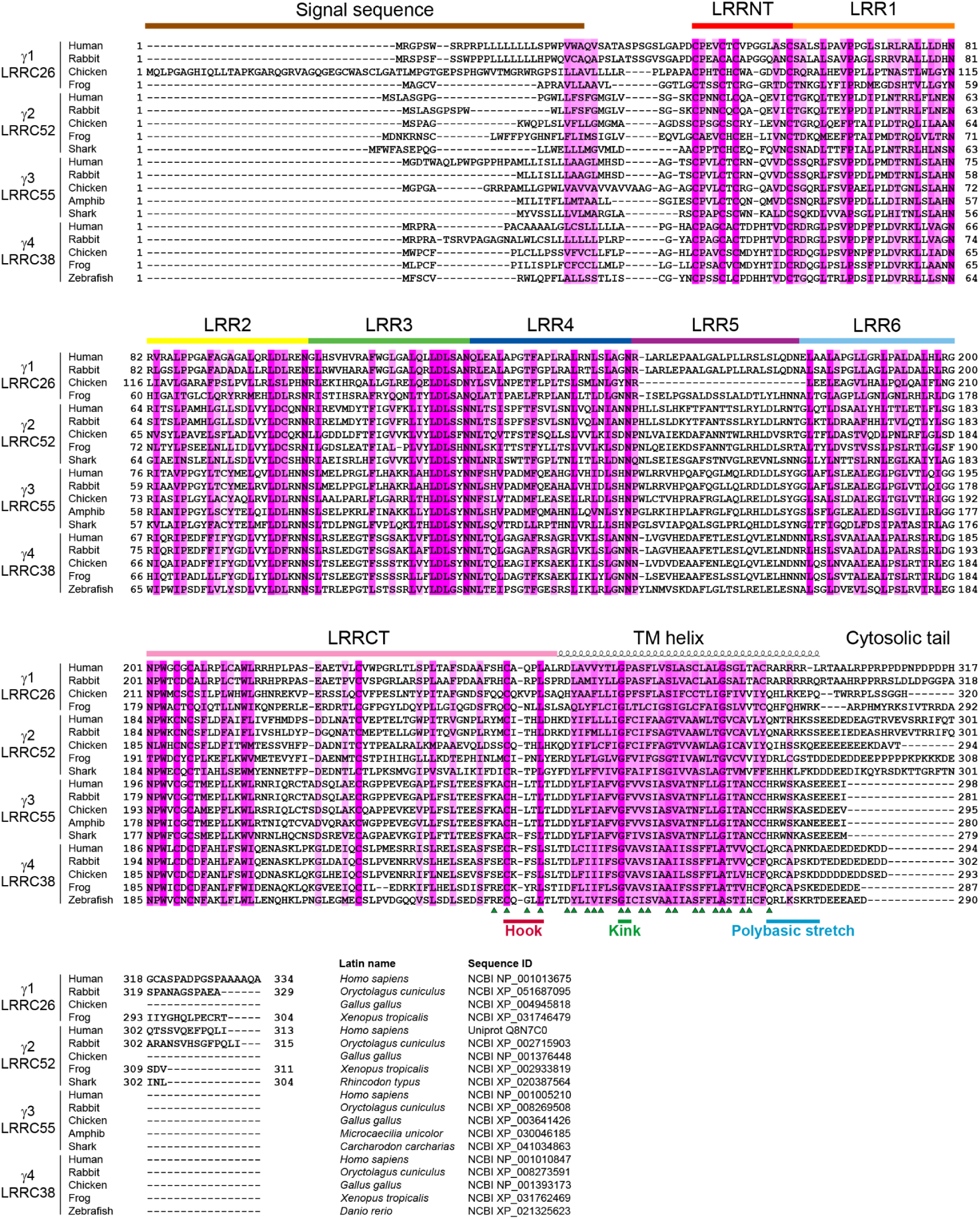
Sequence alignment of γ subunits. Multi-sequence alignment of the four γ subunit families from different vertebrate species. Species names and accession numbers are shown in the bottom of the figure. Positional conservation is indicated by the strength of pink background color. Structural elements are indicated above the alignment and residues interacting with Slo1 by green triangles below the alignment.

**Supplementary Figure 2:**
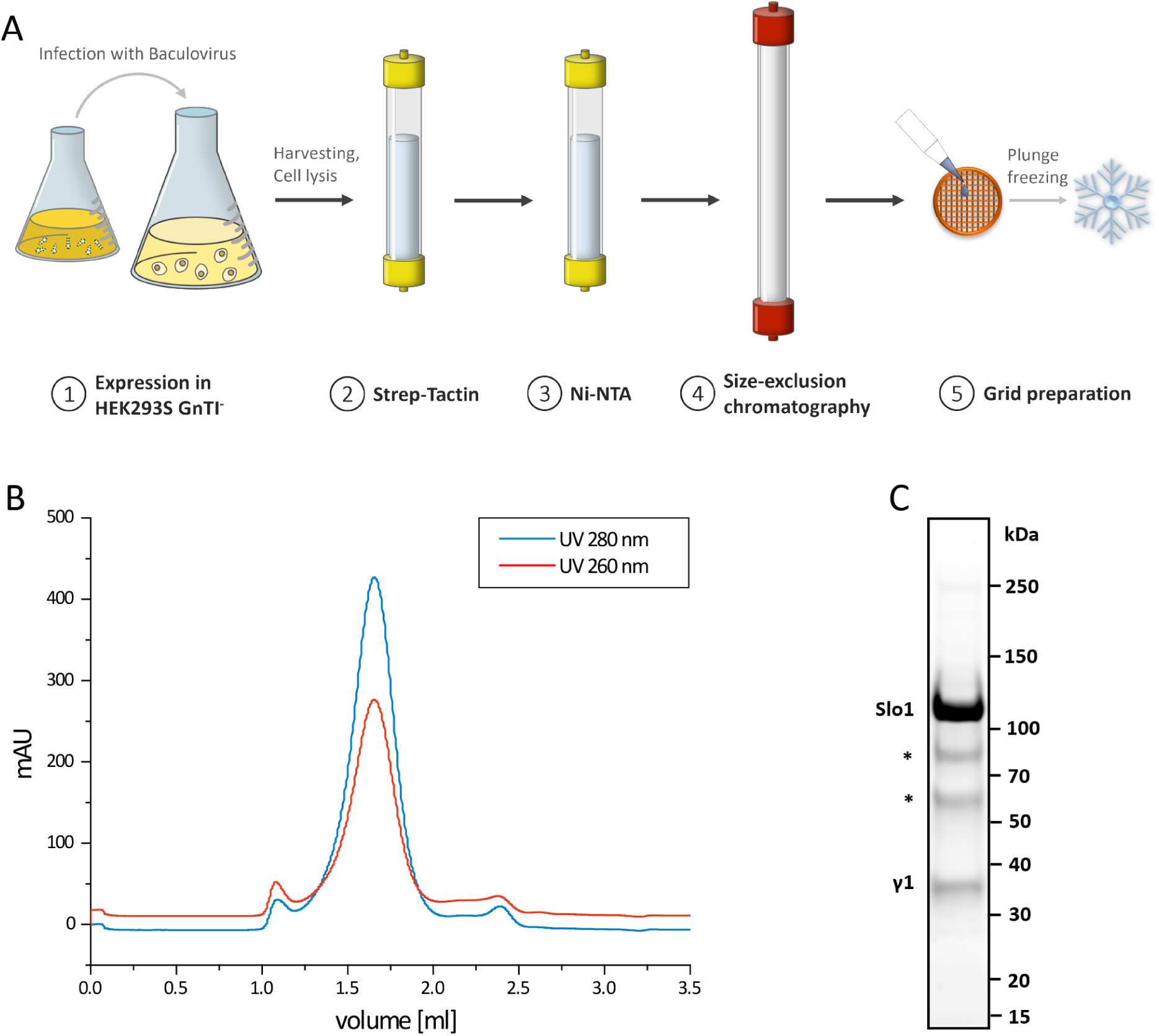
Purification of the Slo1-γ1 complex. A) Purification scheme of Slo1-γ1. The complex was co-expressed by co-infecting HEK293S GnTI- cells with modified baculoviruses containing genetic information for Slo1-mCherry-2xStrep and γ1-eGFP-His_10_, respectively. Cells were lysed and the solubilized membranes incubated with Strep-Tactin beads to pull out Strep-tagged Slo1 and co-assembled γ1. The eluate was further purified via Ni-NTA resin (to pull on His_10_-tagged γ1 and co-assembled Slo1). The double affinity-purified sample was incubated overnight with HRV-3C protease to remove the fluorescence tags and then applied to size exclusion chromatography via Superose 6 5/150. The purified, concentrated protein was then subjected to cryo-EM grid preparation and cryo-plunging in liquid ethane. B) Size exclusion profile of the Slo1-γ1 complex. C) SDS-PAGE of purified Slo1-γ1 complex after size exclusion chromatography. Two notorious degradation products of Slo1 (marked by asterisks) were always there and represent species which were proteolytically cleaved in an extended linker region in the gating ring.

**Supplementary Figure 3:**
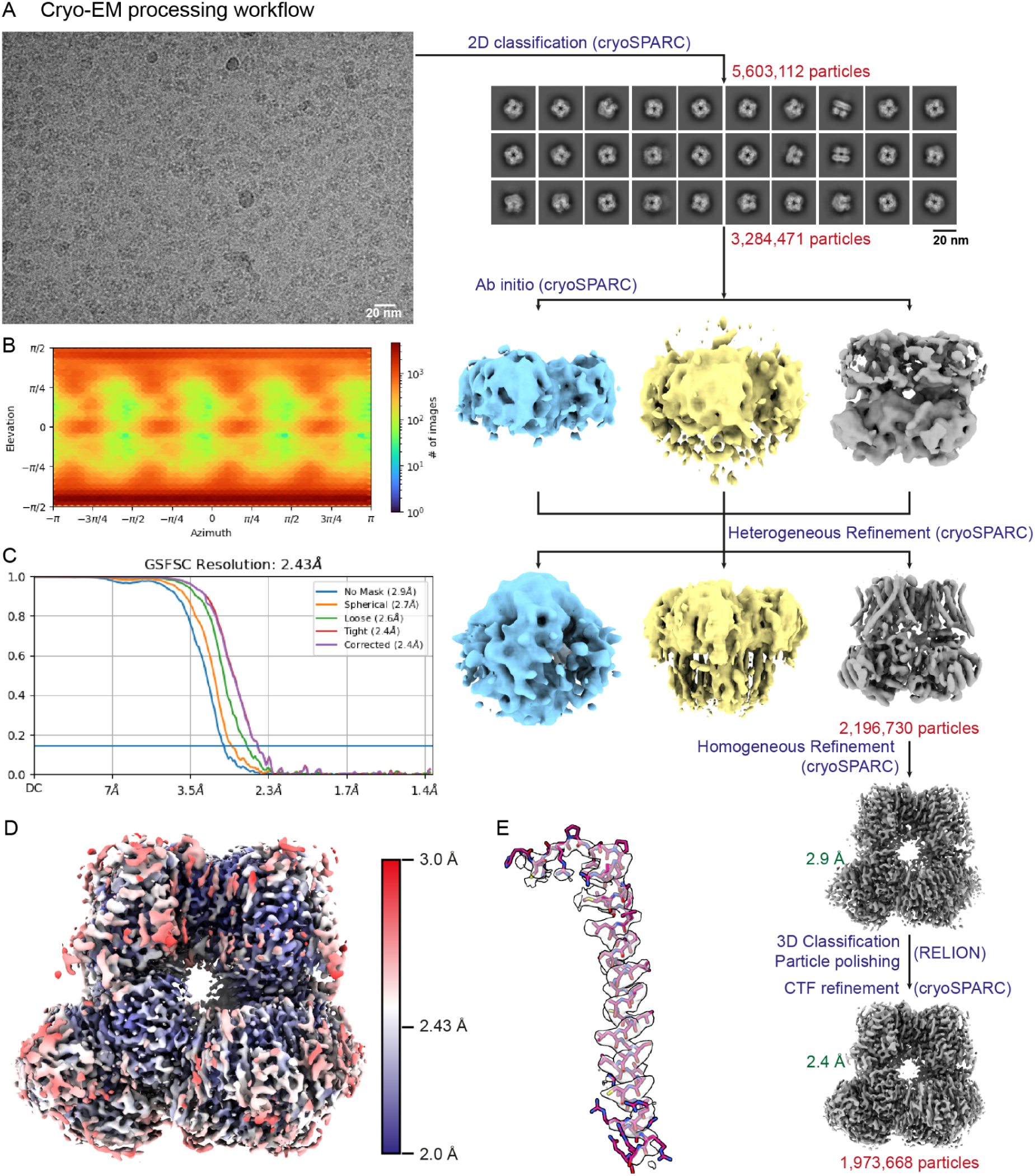
Cryo-EM data processing of the Slo1-γ1 complex. A) Processing workflow including a representative micrograph, 2D classes and intermediate and final reconstructions. The final reconstruction reached 2.43 Å and contained 1,973,668 of the 5,603,112 initial particles. B) Particle orientation distribution plot. C) Gold-standard FSC curve. The blue horizontal line marks the gold-standard 0.143 threshold. D) Final, locally filtered reconstruction colored by local resolution in a gradient from blue (2.0 Å) over white to red (3.0 Å). E) Quality of the cryo-EM reconstruction of the γ1 transmembrane helix.

**Supplementary Figure 4:**
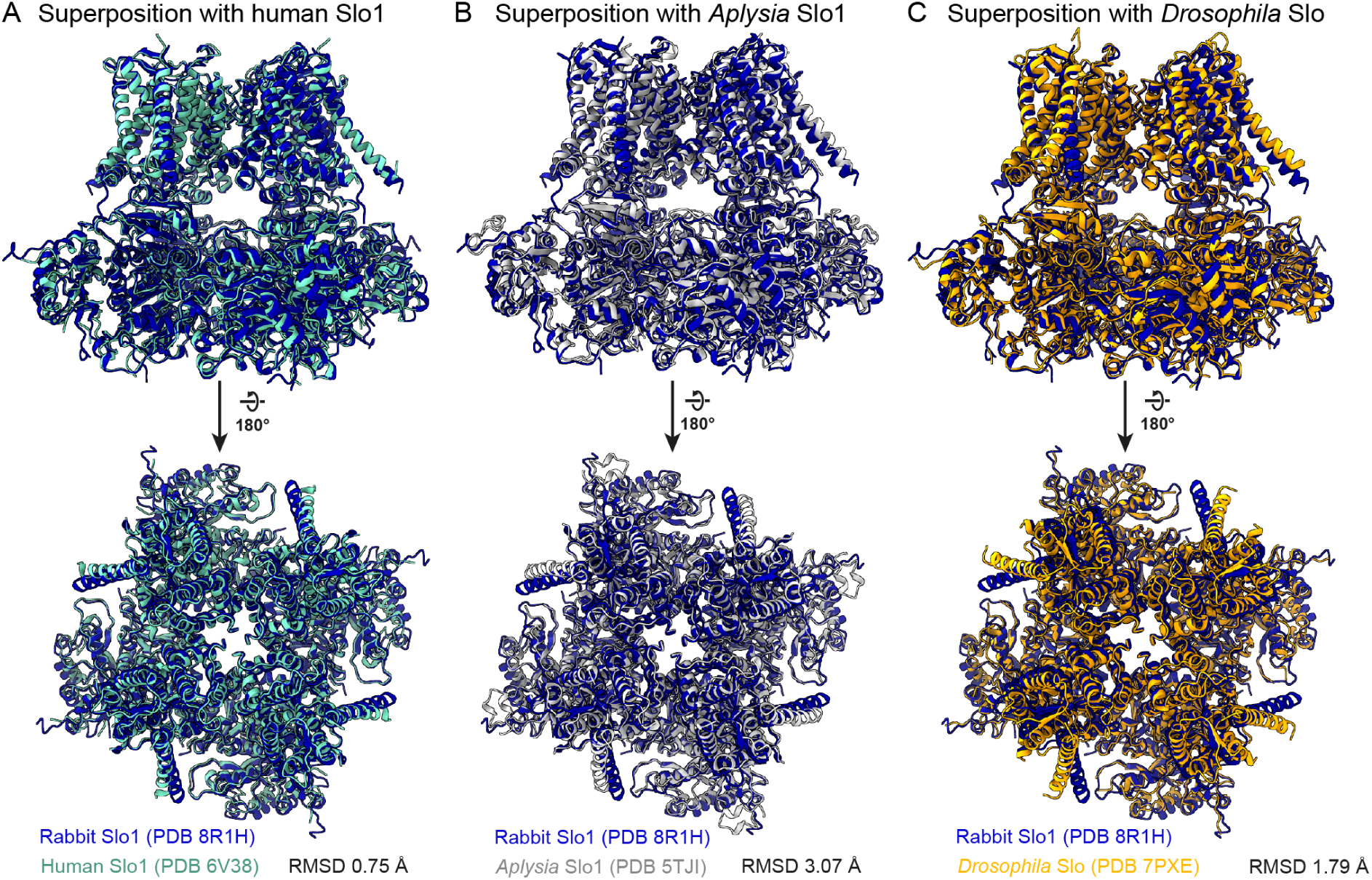
γ1-bound Slo1 adopts a conformation similar to other Slo channel structures. A) Superposition of rabbit Slo1 (blue, this work) with human Slo1 (cyan)^10^. B) Superposition of rabbit Slo1 (blue, this work) with Aplysia Slo1 (grey)^11^. C) Superposition of rabbit Slo1 (blue, this work) with Drosophila Slo (gold)^8^.

**Supplementary Figure 5:**
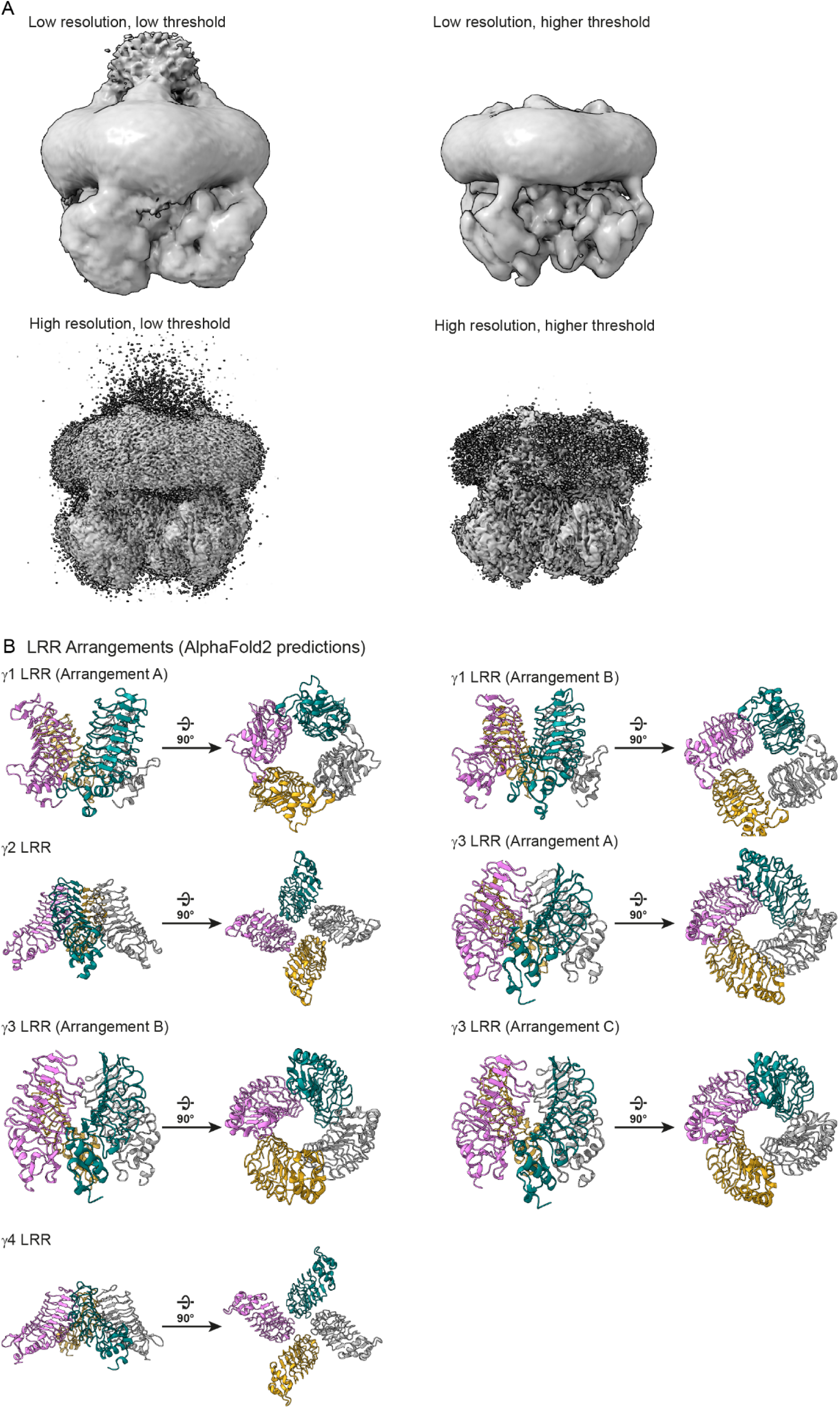
γ1 LRR domains likely assemble as tetramers to form a flexible extracellular module. A) Cryo-EM reconstruction filtered at 6 Å (top row) and at 2.43 Å (bottom row), shown at low (left) and higher (right) threshold. A diffuse density is visible in the low-threshold panels, indicating the presence of a weakly ordered extracellular module above the transmembrane module. B) AlphaFold-Multimer^31^ predictions of γ subunit LRR domains. For each γ subunit, 10 models were created. Here, all unique solutions are shown which are structurally compatible with assembly with Slo1, i.e. all four C-termini point in the same direction.

## Notes

### Competing Interest Statement

The authors have declared no competing interest.

